# A machine-vision approach for automated pain measurement at millisecond timescales

**DOI:** 10.1101/2020.02.18.955070

**Authors:** Jessica Jones, William Foster, Colin Twomey, Justin Burdge, Osama Ahmed, Jessica A. Wojick, Gregory Corder, Joshua B. Plotkin, Ishmail Abdus-Saboor

## Abstract

Objective and automatic measurement of pain in mice remains a barrier for discovery in both basic and translational neuroscience. Here we capture rapid paw kinematics during pain behavior in mice with high-speed videography and automated paw tracking with machine and deep learning approaches. Our statistical software platform, PAWS (Pain Assessment at Withdrawal Speeds), uses a univariate projection of paw position over time to automatically quantify fast paw dynamics at the onset of paw withdrawal and also lingering pain-related behaviors such as paw guarding and shaking. Applied to innocuous and noxious stimuli across six inbred mouse strains, a linear discriminant analysis reveals a two-dimensional subspace that separates painful from non-painful stimuli on one axis, and further distinguishes the severity of pain on the second axis. Automated paw tracking combined with PAWS reveals behaviorally-divergent mouse strains that display hypo- and hyper-sensitivity to mechanical stimuli. To demonstrate the efficacy of PAWS for detecting hypersensitivity to noxious stimuli, we chemogenetically activated pain-aversion neurons in the amygdala, which further separated the behavioral representation of pain-related behaviors along a low-dimensional path. Taken together, this automated pain quantification approach should increase the ease and objectivity of collecting rigorous behavioral data, and it is compatible with other neural circuit dissection tools for determining the mouse pain state.

## INTRODUCTION

Numerous genetic and environmental factors shape the subjective experience of pain. While humans can articulate the intensity and unpleasantness of their perceived pain in the form of pain scales and questionnaires[1, 2], determining pain states in non-verbal animals remains a significant challenge. Rodents are the predominant model organism to study pain and there is an urgent need to develop high-throughput approaches that accurately measure pain. The past fifty years of pain research have relied on the paw withdrawal reflex metric to measure pain-related behaviors in rodents, which has contributed to important discoveries about nociception[3, 4]. However, the traditional approach of manually scoring paw lifting suffers from an inability to determine whether paw movement away from a stimulus is motivated by the experience of pain. Improving the resolution, and increasing the dimensionality, of the common paw withdrawal assay has the potential to increase the predictive validity of translational pain therapeutics, and to increase the rate at which basic science findings are translated to the clinic.

Animals generate rapid motor responses to somatosensory stimuli at millisecond speeds that cannot be readily detected by eye[5, 6]. Therefore, significantly increasing the recording rate of the motor actions, coupled with sub-second mapping of behavioral signatures, will sharpen the resolution and confidence for assessing an animal’s internal pain state. For example, researchers recorded optogenetically-induced nociceptive behaviors at 240 frames per second (fps), which facilitated the precise mapping of nocifensive behaviors including paw withdrawal, paw guarding, jumping, and vocalization [7]. Two additional studies recording between 500-1,000 fps using both natural and optogenetic nociceptive stimuli, demonstrated that nociceptive withdrawal latencies were on the order of 20-130 milliseconds (ms)[8, 9]. More recently, we recorded mouse somatosensory behaviors at 500-1,000 fps, coupled with manual behavioral mapping, statistical modeling, and machine learning to create a more objective “pain scale” [10]. Although these studies provide a framework for using high-speed videography for fine-assessment of pain, a major limitation lies in the relatively low-throughput nature of manual scoring of the video frames, which adds potential human error, and limits the ease of platform adoption in other laboratories.

Recently, computational neuroethology platforms have introduced a suite of machine learning and deep neural networks to automatically track animal body parts during behavior for postural estimation [11]. Platforms such as Motion Sequencing (MoSeq) use 3-dimensional depth imaging, quantitative analyses, and fitting with unsupervised computational models to estimate animal posture within an open arena and can automatically reveal ∼60 unique sub-second behavioral signatures [12]. DeepLabCut and LEAP, train deep neural networks (DNN) with relatively limited training datasets, allowing the computer to accurately track unlabeled body parts such as a mouse paw, ear, or even a single digit through many frames of videography data [14, 15]. Alternatively, the markerless automated tracking software ProAnalyst, tracks moving objects across high frame rate videography data [16, 17]. This approach does not use deep learning but relies on built-in machine learning algorithms for automated tracking, which provides an easier point of entry for researchers with limited time for software development or computing power.

Here, we present an automated mouse pain scale that combines videography at 2,000 fps, automated paw tracking with ProAnalyst and Social LEAP, and new software called PAWS (Pain Assessment at Withdrawal Speeds), which scores eight defined behavioral endpoints. Beginning with six commonly used genetically inbred mouse strains we revealed stereotyped sub-second paw trajectory patterns, with simple up-down lifts typifying the response to innocuous stimuli and elaborate sinusoidal patterns typifying the responses to noxious stimuli. By projecting paw position onto the time-varying principal axis of paw movement, we identified shaking behavior as simple sequences of peaks and valleys in this univariate time series, and paw guarding as extended periods of stasis devoid of shaking prior to returning the paw to the ground.

After building an automated pain assessment platform, we used statistical modeling with linear discriminant analyses (LDA) and confirmed that the eight movement features we automatically measured were sufficient to separate behavioral responses to innocuous touch from noxious pinprick stimuli. The LDA further segregated noxious intensity. Cross-validation of our LDA modeling confirmed that PAWS performed better than an unsupervised machine learning approach for decoding the stimulus type and intensity based on the animal’s sub-second behavioral responses. Lastly, using our recently described protocol to gain genetic-access to basolateral amygdala (BLA) neurons that are responsive to pain [19], we chemogenetically activated the BLA pain ensemble to validate our platform’s ability to detect pain hypersensitivity. Thus, we can automatically measure increases in mechanical pain responsiveness to noxious stimuli while manipulating central pain circuits. Taken together, this work reveals that automating paw tracking and subsequent quantification of pain behaviors with high frame rate videography provides a reliable method to objectively determine the mouse pain state.

## RESULTS

### High-speed videography and automated paw tracking during evoked behaviors

In order to capture sub-second behavioral ethograms during somatosensory behaviors in freely behaving mice, we recorded mice at 2,000 fps, with a particular focus on the stimulated paw. We reasoned that we could develop a pipeline where we first performed behavior experiments, followed by automated paw tracking and automated pain scoring, and lastly statistical modeling to transform multidimensional datasets into a single dimension that separated touch from pain (Figure 1A). To begin, we used 10 mice from six commonly used inbred lines and on separate days to avoid sensitization to the stimuli, we applied to one hind paw a static innocuous stimulus (cotton swab), a moving innocuous stimulus (dynamic brush), a weak noxious stimulus (light application of a pinprick), and an intense noxious stimulus (heavy application of a pinprick). Using traditional pain scoring, we noticed that the paw withdrawal frequencies to these four stimuli varied widely across these six strains. For example, Balb/cJ, DBA1/J, and CBA/J displayed high paw withdrawal rates to all mechanical stimuli, both the innocuous and the two noxious (Figure 1B). Conversely, C57BL/6J and AKR/J displayed high paw withdrawal rates to both the noxious stimuli and the innocuous dynamic brush, but not the cotton swab (Figure 1B). Lastly, the A/J mice displayed high rates of paw lifting to both pinprick stimuli and low withdrawal rates to the two touch stimuli (Figure 1B). Since mice will move their paw to both innocuous and noxious stimuli, it is hard to determine if these differences in withdrawal frequencies are driven by genetic differences in susceptibility to pain. What these data likely reveal using the common traditional approach is that this test in isolation may be an inadequate measurement of pain at baseline states.

**Figure 1.**
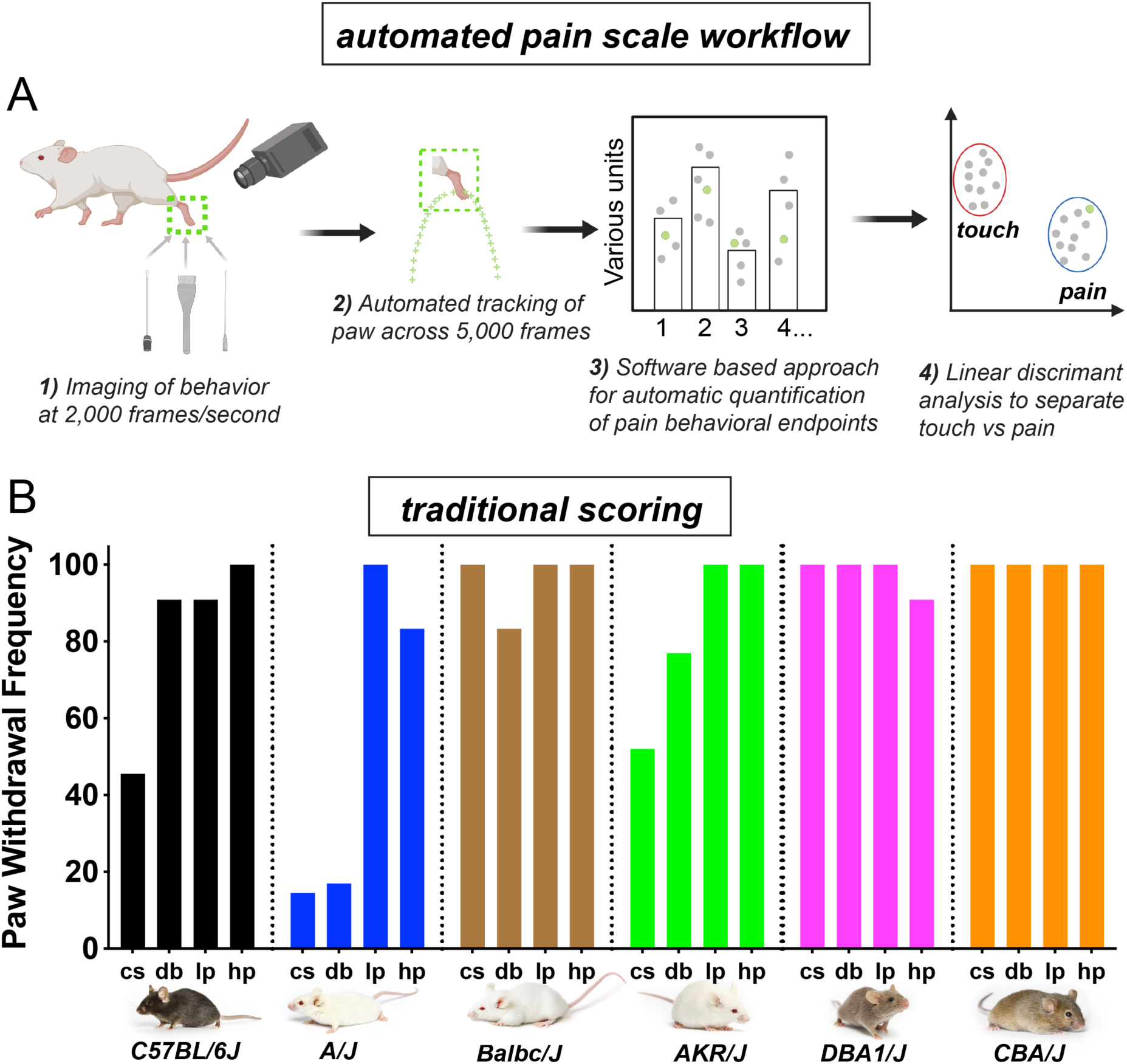
Automated pain assessment workflow in comparison to traditional unidimensional pain scoring. **a**, Workflow pipeline in following order consisting of: 1) high-speed videography of freely behaving mice, 2) machine/deep learning based approaches for automatic tracking of the stimulated paw, 3) PAWS software for automatic quantification of defined pain behavioral endpoints, 4) statistical modeling with LDA for separation of touch versus pain on trial-by-trial basis. **b**, Traditional scoring focused on paw withdrawal frequencies to four mechanical stimuli: cs=cotton swab, db=dynamic brush, lp=light pinprick, hp=heavy pinprick. N=10 mice per strain. Images from Jackson laboratories.

Next we turned to automated tracking with the six mouse strains to determine the X,Y coordinates of the paw across approximately 5,000 frames – recording at 2,000 fps with total behavior time from stimulus application to paw lift and return back to the floor being approximately 2-3 seconds. Using machine learning algorithms embedded within the ProAnalyst motion tracking software, we manually labeled the center of the stimulated paw in each video and the machine automatically tracked the paw throughout each additional frame (Figure 2A-X) (see Movie 1). While observing the automated paw trajectory patterns we noticed that stereotyped motor sequences defined the movement away from the four stimuli, regardless of strain background (Figure 2A-X). For example, the responses to the two innocuous stimuli were typically up-down C-shaped movements (Figure 2A-X). Conversely, the responses to the two noxious stimuli were typically more elaborate movements, often accompanied by orbital tightening of the eye, which is a known facial feature of intense pain (Figure 2A-X) [20]. In the majority of tested strains, we also noticed that the paw trajectory pattern in response to a weakly painful stimulus (light pinprick) often resulted in a figure-eight like sequence that may point towards a shared sensorimotor neural circuit that governs how animals respond to weakly painful stimuli given their body posture and space constraints (Figure 2A-X).

**Figure 2.**
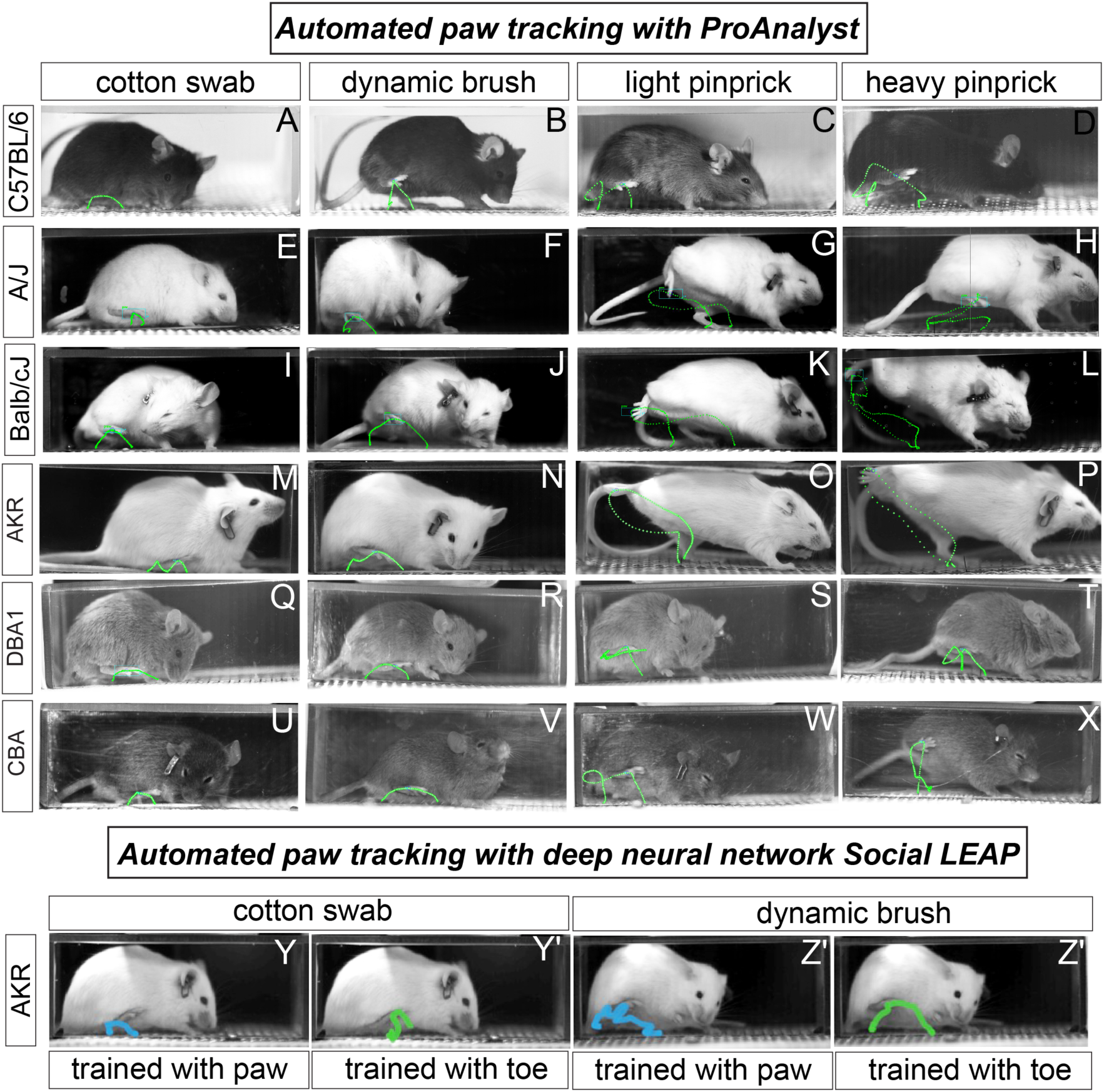
Automated paw tracking with high-speed recording of behavior. **a-x**, ProAnalyst machine learning based paw tracking. **y-z′**, Social LEAP deep neural network based paw or toe tracking. All still images represent a single frame of ∼5,000 total frames. Green and blue lines display paw trajectory patterns during the entire behavior. N=10/mice per stimulus and images shown are representative of each strain.

Next, using the AKR/J strain we used a deep learning-based pose tracking algorithm called Social LEAP (in preparation, based on[14]), to predict mouse toe and mid-paw positions during somatosensory behaviors recorded at high speed (Figure 2Y-Z′). Our mouse paw-tracking model was generated from a small training set of video frames collected from the four assays (∼9.5% of video frames per assay). In general, the paw trajectory patterns with Social LEAP resemble those of ProAnalyst, and the software package we describe below is compatible with automated tracking data from either tool. Taken together, our ability to detect clear qualitative distinctions in paw movements with automated tracking approaches, gave us confidence that we could use spatiotemporal data of paw position to automatically extract features that may be useful in determining the mouse pain state.

### Development of software to automatically score pain behavioral endpoints

We developed software to systematically quantify seven pain-relevant features of the paw position time series based on existing measurements in the literature. Maximum paw height, lateral velocity, vertical velocity, and the total distance traveled by the paw, were all computed based on a polynomial smoothing (Savitsky-Golay filter of order 3) of the original paw position time series. These four features were computed for two different windows of the paw trajectory time series: the time leading up to the initial paw peak (time t* in Figure 3A), and the time after the initial paw peak, which we refer to as pre-peak and post-peak, respectively. In the post-peak time window we additionally identified periods of “shaking” and “guarding” based on a threshold displacement along the principal axis of paw movement (Figure 3B and C). We used this delineation to quantify the total duration spent shaking or guarding as well as the total number of paw shakes across shaking periods. We refer to this software package as PAWS (Pain Assessment at Withdrawal Speeds).

**Figure 3.**
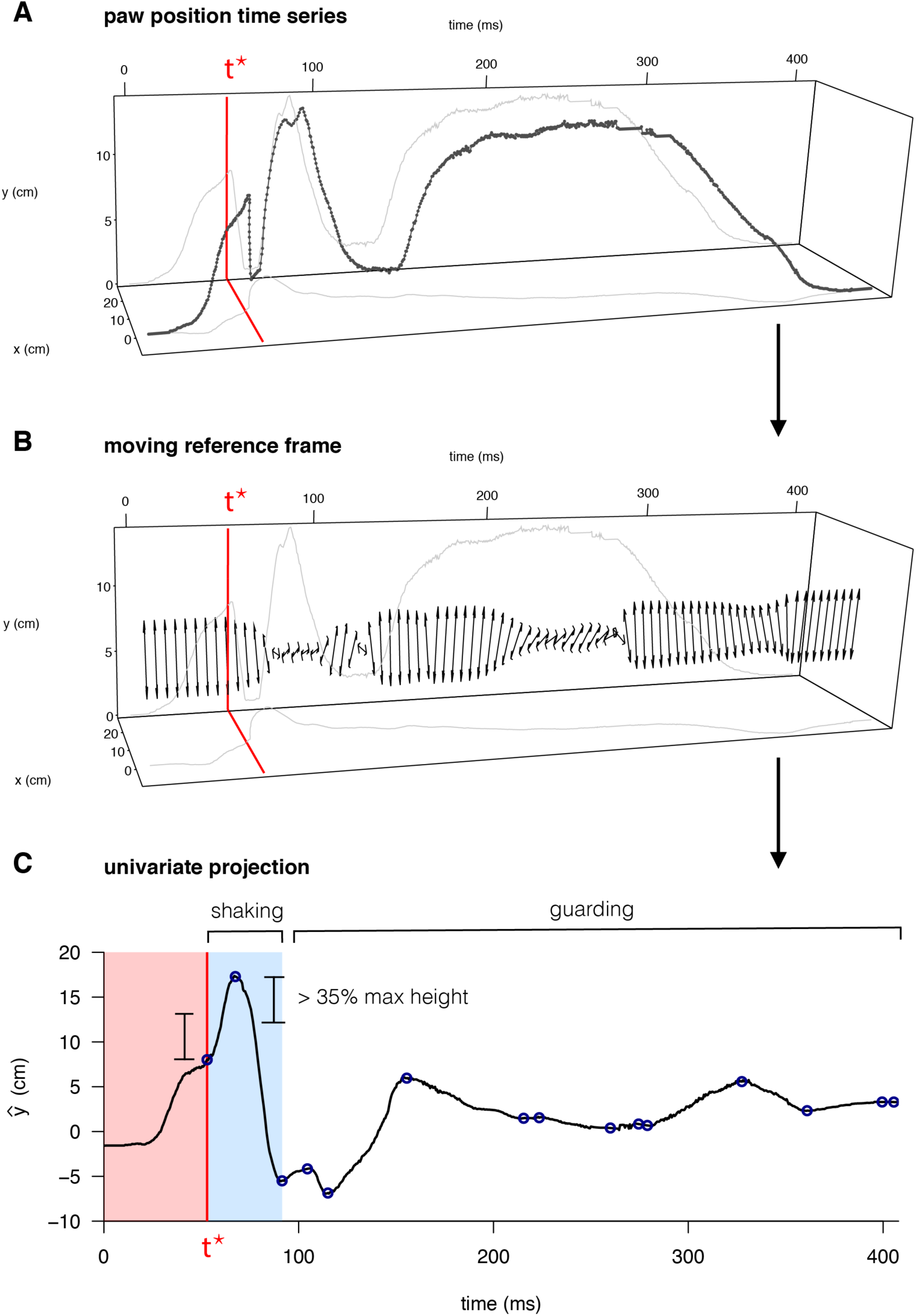
Quantification of behavioral features for mouse pain state. **a**, Raw paw positions (x,y) measured in the camera reference frame (anterior/posterior displacement and vertical height, respectively) as a function of time, with the time of first peak in paw height, t*, marked in red. **b**, The principal axis of paw displacement in a moving time window, shown as a function of time. **c**, Displacement in the principal axis moving reference frame was used to quantify instances of paw shaking in terms of sequences of displacements above a given threshold relative to the maximum paw height displacement.

### Automated scoring of rapid paw dynamics and lingering pain behaviors

With software generated to automatically measure paw movement features related to the mouse pain state, we plotted and analyzed the data across the six mouse strains with the four mechanical stimuli described above. For plotting the individual behavioral endpoints, we separated the paw distance traveled measurement into pre-peak and post-peak distances. To standardize across varying units and to appreciate individual deviation from the median, we transformed the raw output measurements into Z-scores (Figure 4). The first readily apparent feature we noticed was that all measurements across these six strains were interspersed without clear separation amongst strains, despite the vast differences in paw withdrawal frequencies to these same stimuli (Figure 1B). In regards to pre paw peak behavioral endpoints, we observed a statistical separation in the stimulated paw’s velocity on both the X- and Y-axes between touch (CS,DB) and pain stimuli (LP,HP) as the paw withdrew upwards to reach its first highest peak (Figure 4A-D). Relatedly, the max height that the stimulated paw reached its first highest peak, as well as the distance the stimulated paw traveled to reach that peak, separated out statistically comparing the innocuous versus painful stimuli (Figure 4A-D).

**Figure 4.**
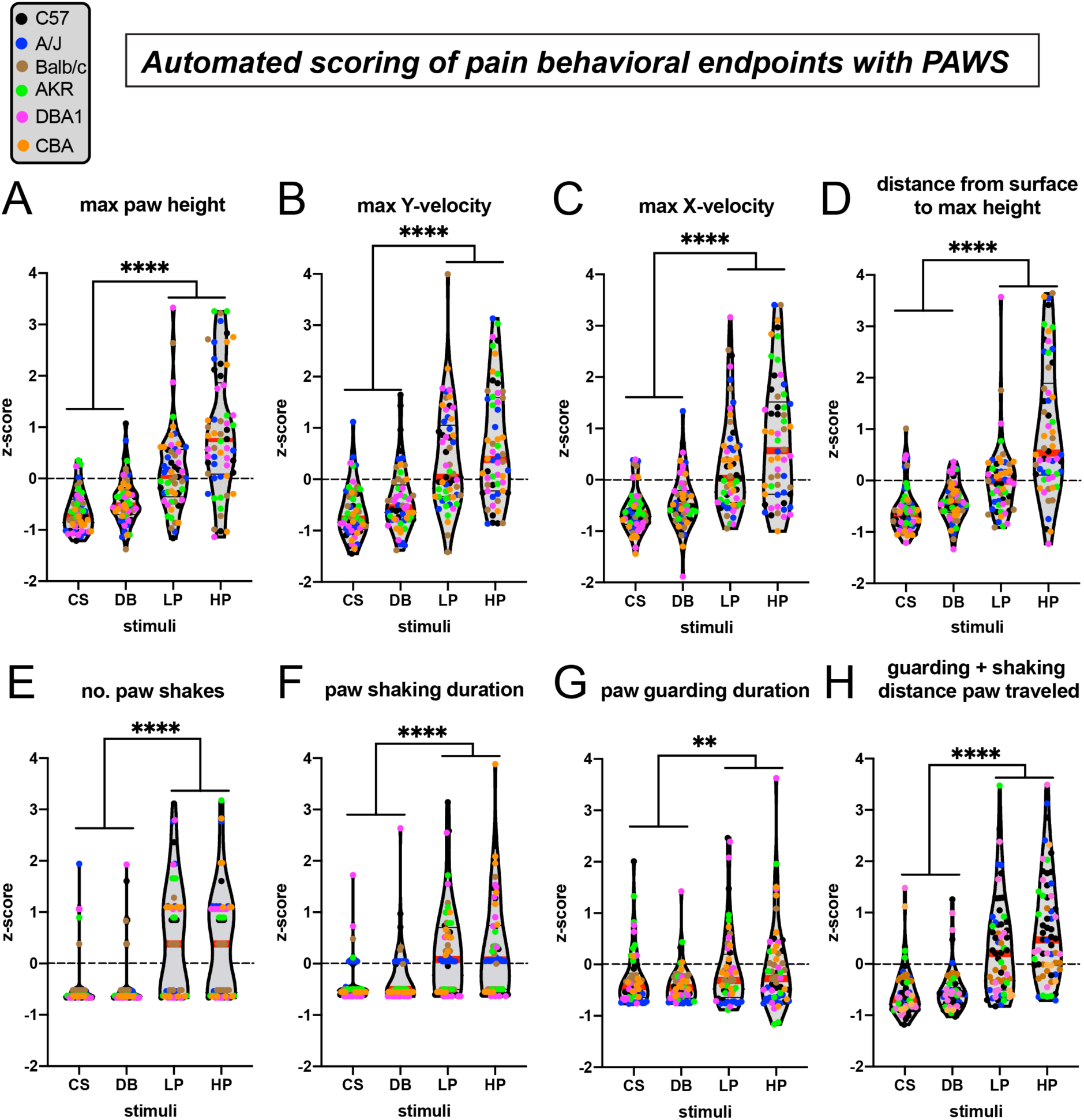
Automated measurement of pain behavioral endpoints across strains with new software PAWS. Measurements are converted to Z-score to reveal deviation from mean of individual measures and to standardize units across the eight measures. **a-d**, Pain measurements of the stimulated paw from lift to max height. **e-h**, Pain measurements of the stimulated paw from max height to paw return. N=10/mice of each strain given each stimulus once. Statistical significance was computed by comparing CS+DB versus LP+HP with Wilcoxon matched-pairs signed rank test. ** represents p-value ≤ 0.01. **** represents p-value ≤ 0.0001. On the violin plots, black horizontal lines represent quartiles and red horizontal lines represent the median.

We next plotted the pain behavioral endpoints that occur after the paw has reached its highest first peak and before the animal places its paw back to the surface. This time window corresponds to a time when the animal engages supra-spinal neural circuits and consciously decides whether to engage in coping behaviors like paw shaking or defensive behaviors like paw guarding. Similar to our observations with the four pre paw peak pain behaviors, the four post paw peak behaviors also show statistical separation between our innocuous and pain stimuli (Figure 4E-D). Although we are not the first research group to observe paw shaking and guarding behaviors in rodents, this is one of the first technologies to automatically score the number and duration of these paw movement dynamics. Taken together, these automated measurements extracted from paw time series data are sufficient to objectively separate touch from pain in genetically diverse mice.

### Statistical modeling with linear discriminant analyses separates touch versus pain across six inbred mouse strains

We used linear discriminant analysis (LDA) to identify a linear subspace of the quantified behavioral features that best separates the four pain treatments: CS, DB, LP, and HP. We did this for two cases: first, restricting to just four features quantifiable pre-peak (t<t*), and then for all seven features quantified post-peak (Fig. 5A and B, respectively). The proportion of the trace accounted for by each of the first two components of the linear discriminant analysis (LD1 and LD2) was 65.6% and 34.2%, respectively, in the pre-peak LDA, and 90.9% and 8.5%, respectively, in the post-peak LDA. In both cases LD3 (not shown) accounted for < 1% of the trace. For both the pre- and post-peak paw features, the two non-pain stimuli (CS and DB) cover the same parts of the subspace and are largely indistinguishable. In contrast, the low and high pain conditions (LP and HP) separate from the non-pain stimuli and also separate from each other in this subspace.

**Figure 5.**
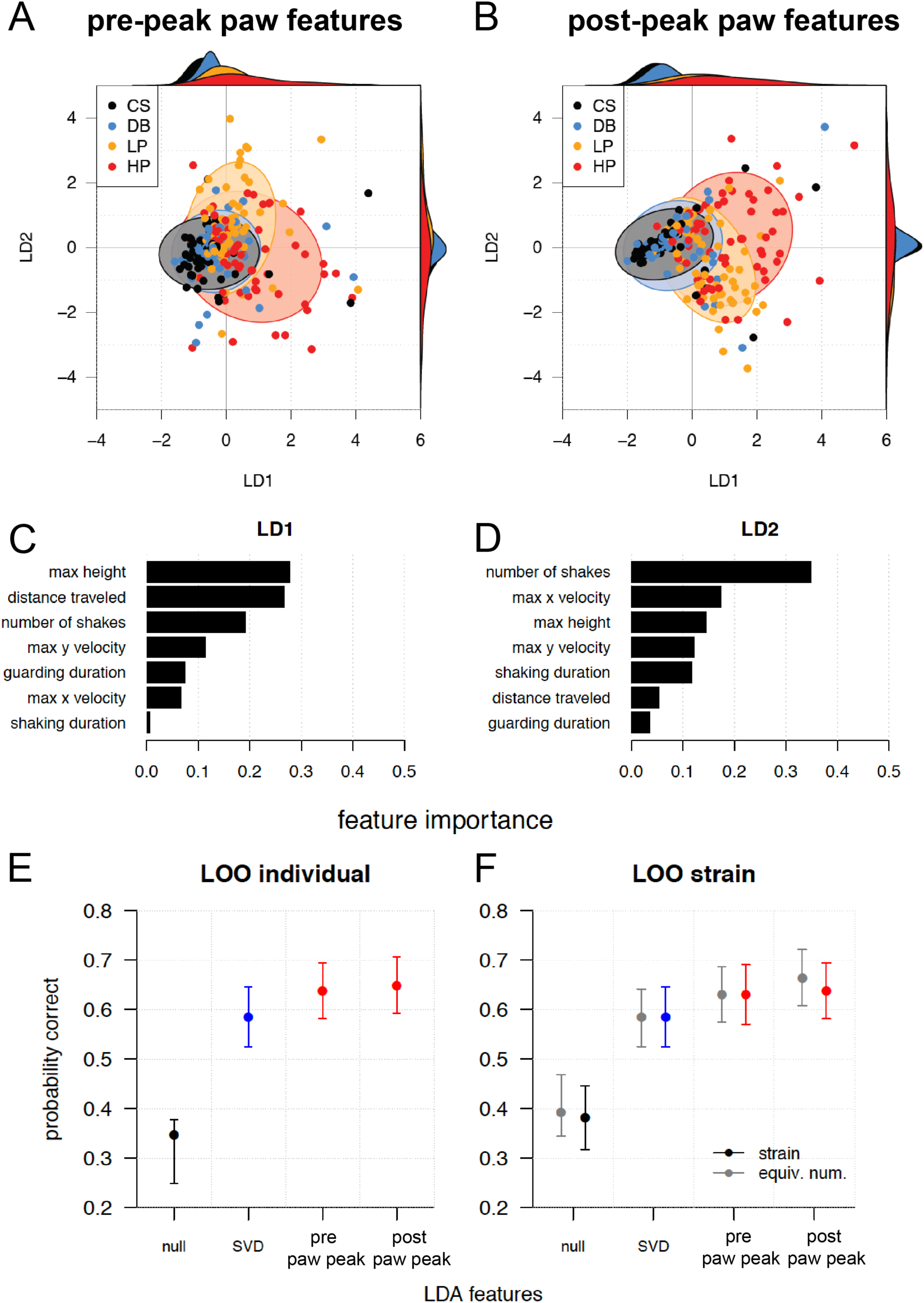
**a**, First two linear discriminant dimensions that best separate the four pain states studied: CS, DB, LP, and HP, for pre-peak features only. Ellipses show Gaussian 95% confidence region for each pain state. Univariate kernel density estimates for LD1 and LD2 decomposed by pain state are shown at top and right, respectively. **b**, Same as **a** but for post-peak paw features only. **c**, Weights assigned to standardized (mean subtracted, variance scaled) behavioral features on the first linear discriminant dimension (LD1) of the post-peak paw features LDA. **d**, Same as **c** but for LD2. **e**, Leave-One-Out (LOO) cross-validation performance of the pre-peak and global LDAs, compared to automatically generated features via SVD, and a null model, for individual mice. **f**, Same as **e**, but leaving out an entire strain (colors) or an equivalent number of mice chosen uniformly at random (gray).

The relative importance of each standardized (mean subtracted, variance scaled) behavioral feature was quantified as the loading of that feature on a given axis of discrimination. The feature importance for the first two linear discriminant dimensions (Figure 5C and D, respectively) indicate that relatively simple properties of the paw trajectory, such as the maximum height of the paw and the total distance traveled before the paw returns to resting position, contribute substantially to separating pain from non-pain (LD1). The number of paw shakes further contributes to the separation of low versus high pain categories in LD2.

Model performance was evaluated by leave-one-out (LOO) cross-validation for individual mice (Figure 5E), and by leave-one-strain-out for strains (Figure 5F). We compared the pre- and post-peak paw LDAs to (1) an LDA based on automatic feature extraction, using 23 features obtained from a generic method to inferring features from variability in the time series themselves (see Methods); and (2) a null model that assigns pain classes according to their probability in the training data, without reference to measured behavioral features. For prediction, CS and DB were treated as a single “non-pain” class, while LP and HP were kept separate, resulting in three classes: non-pain, low pain, and high pain. For predicting the pain state of a given mouse, or all the mice in a strain (generalizing across strains), post-peak paw features consistently performed best. The eight behavioral features we defined and extracted from paw-tracked videography significantly outperform both a random null model as well as the 23 features identified automatically from the same data, in terms of cross-validated ability to discriminate pain stimulus.

### Validation of the automated tracking and scoring approach with new mouse lines

In the process of testing the six inbred lines described above, we tested an additional two strains that appeared to have atypical behavioral responses to our four somatosensory stimuli. The first of these two strains, 129S1, appeared to have a pain hypo-sensitivity phenotype, responding to both innocuous and noxious mechanical stimuli with the same basic up-down paw trajectory pattern, and not the elaborate paw withdrawal pattern typically observed with noxious stimuli (Fig 6A-D) (compare to Figure 2A-X). Prior studies using traditional unidimensional assays to compare the baseline mechanical pain sensitivity of different 129 sub-strains to C57BL/6J mice revealed mixed results with some tests showing no differences between the strains, and either greater or reduced sensitivity to pain [21-23]. The second apparent outlier strain were the SJL mice, where we observe the opposite of what we found in 129S1, where this strain appears hypersensitive to mechanical pain – responding to the soft dynamic brush as if it were pinprick and having even more elaborate paw withdrawal stimuli (Figure 6E-H). The outlier patterns of these two strains were observed when scoring the individual eight pain behavioral endpoints (Table 1). Consistent with this finding, when we perform our LOO correlation testing to determine how accurate our training is from LDA modeling, for both 129 and SJL strains, our confidence in predicting the stimulus the animal received based on its response is low, proving the nature of the outlier phenotypes in these mice (Figure 6I). These data reveal that our automated pain assessment platform is capable of detecting individual strain differences in susceptibility to mechanical pain.

**Figure 6.**
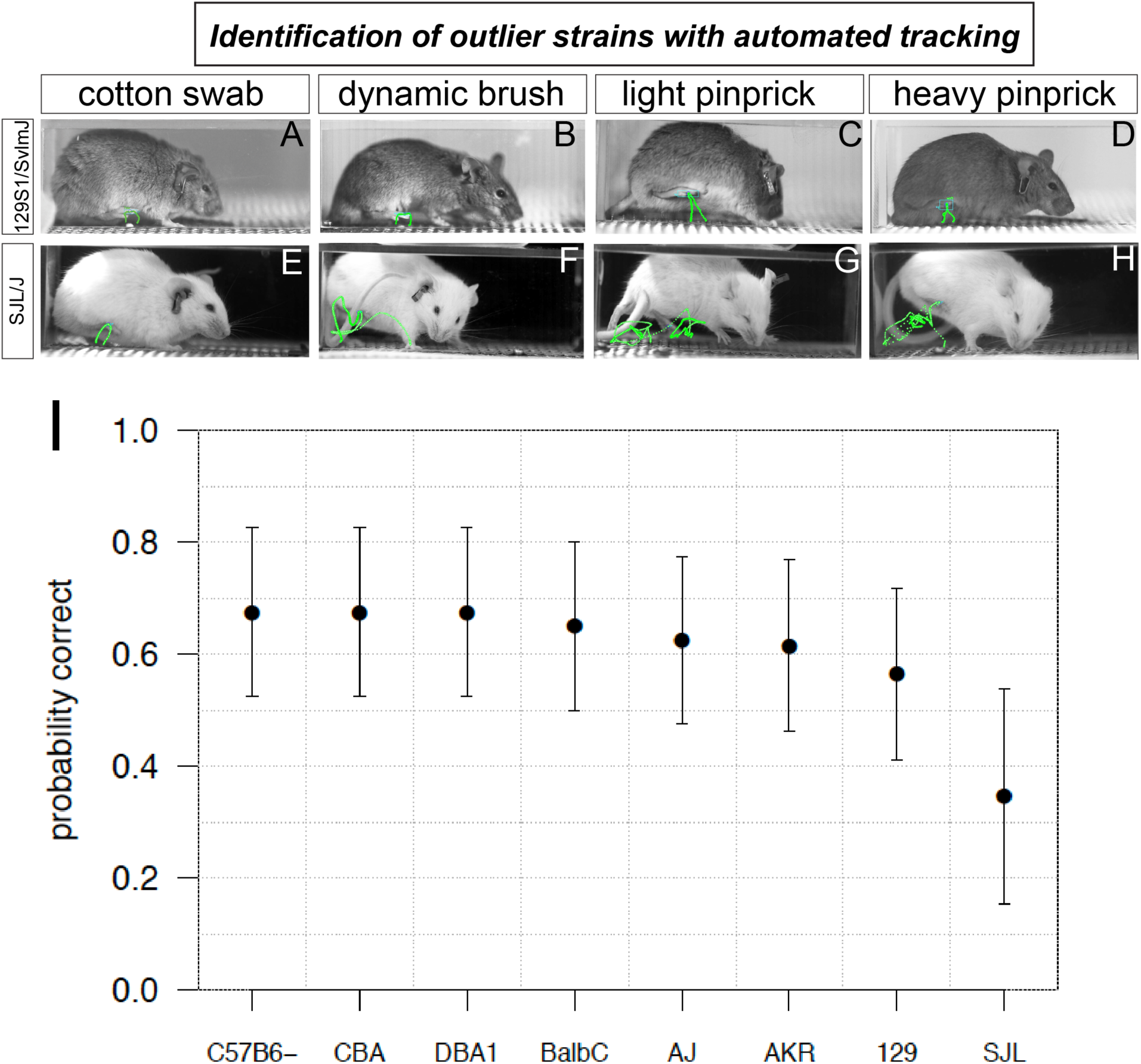
Automated pain assessment platform uncovers two outlier strains. **a-d**, ProAnalyst tracking showing 129S1 mice responding with similar up-down paw lifts to all stimuli, both innocuous and noxious. **e-h**, ProAnalyst tracking showing SJL mice have pain-like response to dynamic brush and heightened pinprick responses. **i**, Leave-out-one (LOO) cross validation performs poorly in predicting the stimulus received in 129 and SJL mice because their responses typically map outside of the normal range of the other 6 strains.

### PAWS reliably detects motivational changes in pain perception following chemogenetic brain circuit manipulation

As another proof-of-principle to validate our ability to automatically measure acute mechanical pain, we next asked if our platform could reliably measure hypersensitivity to the four somatosensory stimuli used above during chemogenetic activation of a neural ensemble encoding pain. To accomplish this, we focused our attention on manipulating pain circuits in the BLA[19]. Briefly, we used the activity-dependent transgenic TRAP2 mice (Fos-FOS-p2A-iCre-ERT2) to gain permanent genetic access to BLA neurons that are responsive to a noxious pin prick to the left hind paw. The transgenic mice in combination with an AAV expressing an excitatory DREADD (AAV5-hSyn-FLEx-hM3q-mCherry) allows the specific expression of the DREADD only in neurons responsive to the noxious pin prick stimulus (*pain*TRAP2^hM3^; Figure 7A). We confirmed bilateral mCherry-labeled DREADD expression in the BLA with this strategy (Figure 7B,C). We first performed our analysis on Cre-negative control animals and observed no effects of CNO injections on behavior. Next, we performed our behavioral analysis on *pain*TRAP2^hM3^ mice at baseline (- CNO) and after activation of the BLA pain ensemble (+CNO) (Figure 7). Because activation of the BLA pain ensemble resulted in unilateral spontaneous guarding pain behaviors, we waited until mice were calm, not moving, and had all four paws on the surface before applying our sensory stimuli. We applied a cotton swab and dynamic brush on one day, and a light and heavy pinprick on a second day. We observed that the PAWS measurements at baseline (-CNO) mirrored those that we observed with the six strains described above, showing separation between innocuous and noxious stimuli. Conversely, with activation of the BLA pain ensemble in *pain*TRAP2^hM3^ mice (+CNO), we noticed increased separation with the post paw peak pain measurements when delivering the heavy pinprick stimuli, most noticeable in the combined shaking/guarding measurement (Figure 7I). When we used LDA for statistical separation that combined our 8 behavioral endpoints, we clearly observe the emergence of a group of mice experiencing a heightened pain state in comparison to the other groups of mice (Figure 7L-O). Thus, PAWS automatically measures increased mechanical pain aversive responsiveness to noxious stimuli during chemogenetic activation of the BLA pain ensemble.

**Figure 7.**
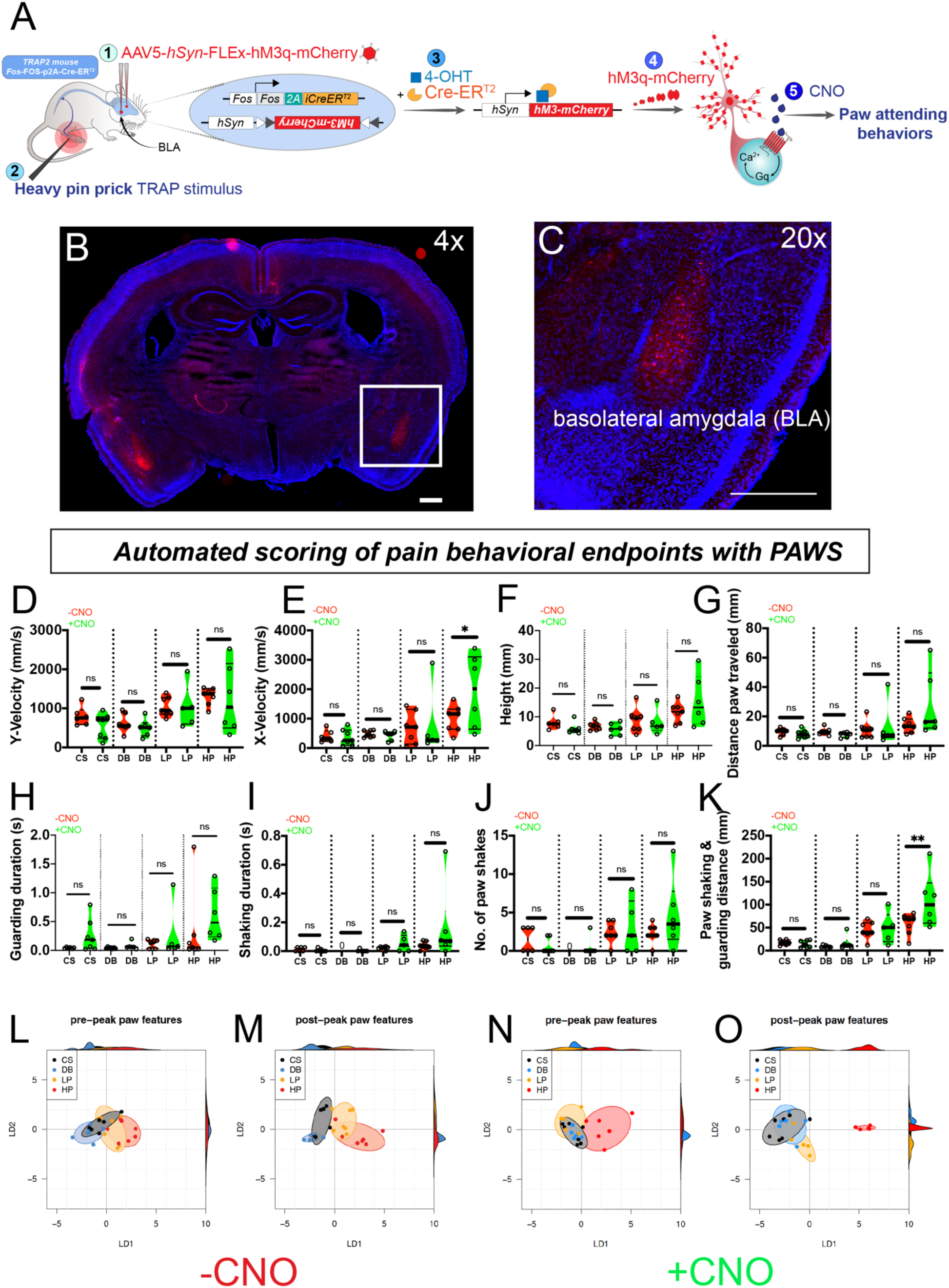
Pain hypersensitivity with chemogenetic activation of the BLA pain ensemble automatically captured via PAWS. **a**, Schematic to permanently tag pain-active neurons in the BLA with an excitatory DREADD in transgenic *pain*TRAP2^hM3^ mice **B.** 4X image of specific and robust expression of m-Cherry-labeled excitatory DREADD bilaterally in the BLA. **c** 20X image of BLA from **b**. Scale bars represent 500μm. **d-g**, Automatic measurement of pre paw peak features comparing mice at baseline (-CNO) to mice administered CNO. (H-K) Automatic measurement of post paw peak features comparing mice at baseline (-CNO) to mice administered CNO. Stimulus abbreviations same as above. Statistical significance was computed with student’s t-test. *represents p-value ≤ 0.05. ** represents p-value ≤ 0.01. **l-o**, Linear discriminant analysis (LDA) using similar factor loadings as described in Fig.5 reveals separation of innocuous vs. noxious stimuli at baseline (-CNO) and a further segregation of the HP group after CNO, only when using all 8 paw features.

## DISCUSSION

Here, we describe an automated approach to quantify the most salient behavioral endpoints following mechanical stimulation of the mouse paw for separating responses according to stimulus intensity. Scoring the paw withdrawal reflex to a natural stimulus is the most commonly used assessment method in pre-clinical rodent models with yes/no responsiveness used as a proxy for inferring pain states. While this methodology for measuring pain in rodents is not fundamentally flawed, it lacks resolution – a limitation that can now be overcome with advances in videography and automated tracking. Here, by automatically tracking paw dynamics at sub-second speeds with high-speed videography coupled with machine learning approaches, we reveal stereotyped trajectory patterns in response to innocuous versus noxious stimuli spanning genetically diverse mice. With an accurate pinpoint of the paw at high spatiotemporal resolution, a freely available software package we term PAWS automatically quantifies eight behavioral endpoints that help to define the mouse pain state. Notably, the eight behavioral features we defined outperform both a random null model as well as 23 unsupervised features extracted from the same data, in terms of their ability to discriminate pain stimuli. Thus, although machine learning is critical for the intermediate task of paw tracking, supervised feature definition still outperforms unsupervised learning for discriminating pain stimuli, and it has the benefit of quantifying pain response in terms of intuitive behaviors (such as paw shakes, guarding, etc). Demonstrating the robustness of the platform, we identify two outlier mouse strains that display reduced or heightened pain sensitivity, and we accurately measure a heightened pain state when we simultaneously activate pain in the periphery and brain using chemogenetic and natural stimuli.

Behavioral neuroscience in model organisms is undergoing a renaissance with the emergence of new tools to automatically track and measure behavior [11]. This renaissance is coincident with many in the research community questioning the robustness of rodent models of pain, addiction, depression, anxiety, and other neuropsychiatric disorders. In kind, many researchers are taking steps backwards to first properly understand the component parts of a complex behavioral sequence before proceeding to identify the neuronal correlates that drive those motor patterns. Both supervised and unsupervised machine learning algorithms are now able to follow unlabeled individual limbs on an experimental animal and automatically define behaviors of interest [12, 14, 15, 24-26]. Although the majority of these tools have yet to be adopted in mass by the pain research community, some of this technology is already in use by pain researchers. For example, the automated grimace scale developed by the Mogil and Zylka labs uses a convolutional neural network trained with 6,000 facial images of mice in “pain” or “non-pain”, to make accurate predictions of the mouse pain state in novel datasets [27]. The automated grimace scale still requires additional customization to assess rodents of different coat colors and to measure chronic pain. Tools like the automated grimace scale that focus on the face, could be combined with the automated pain assessment platform described here that focuses on the paw, for a comprehensive picture of both evoked and spontaneous behavioral responsiveness.

Here, with our platform we observed both homogeneity in behavioral responses across six genetically distinct mouse lines, as well as two outlier strains with responses that mapped outside the range of those six. A wealth of prior literature demonstrated that individual differences in responsiveness to pain in both mice and humans, are driven in part by allelic variation in genes important for pain processing [28, 29]. In regards to the mouse, studies carried out twenty years ago testing pain sensitivity across 11 inbred lines using 12 behavioral read-outs, revealed that depending upon the sensory modality tested, and whether the test was performed before or after injury to the somatosensory system, genotype appeared to influence mouse pain behaviors [30]. A meta-analysis from ten years ago described over 400 papers from mouse pain research that implicated ∼350 genes in pain and analgesia [31]. However, with the relatively limited resolution of some conventional pain behavior assays, it remains unclear how reliable some of these studies are and which potential target genes merit further development as novel analgesics. This assertion is underscored by the fact that only a handful of targets that have shown promise in rodents, have made it to the clinic as novel therapeutics, causing many to question the robustness of the animal models used in pain testing [32, 33]. Here, we uncover a pain hypo-sensitive phenotype in 129S1 mice where the animals respond to pinprick stimuli similar to a soft brush or cotton swab. This finding could have major implications on transgenic mouse lines built using embryonic stem cells from 129S1 mice, especially if sufficient backcrossing to C57BL/6J is not performed. In other words, a pain phenotype may be mistakenly attributed to knocking out a specific gene, while the observed results may be the result of contamination by lingering DNA from the 129S1 donor strain.

Conversely, we observe a phenotype in the opposite direction, where SJL mice respond to a soft brush as if it were a pinprick. To the best of our knowledge, this is the first report of a pain hyper-sensitivity phenotype for SJL mice at baseline. Of note, SJL mice are known to be an aggressor mouse line and even in our studies some animals had to be removed from testing due to excessive fighting between cage mates [34]. Therefore, future studies are necessary to determine if the genetic repertoire that makes these mice aggressive also contributes to their heightened pain responses. Together, with a more precise tool to measure acute mechanical pain at baseline, we can begin to use this platform as a behavioral screening tool followed by subsequent genetic mapping approaches.

As a proof-of-principle to test the robustness of our platform, we demonstrate that we can chemogenetically activate the BLA pain ensemble and detect hypersensitivity to peripheral stimuli. In addition to confirming the precision of our technology, these experiments raise an intriguing biological question: what is the emotional and sensory experience of activating pain-responsive neurons without a peripheral injury? Would tonic activation of this BLA pain ensemble be comparable to human experiences of chronic pain? Further studies to examine these questions are ongoing.

In summary, we have developed a rapid and user-friendly automated pain assessment platform for measuring mechanical pain in mice. Since mechanical stimulation of the rodent hind paw remains the most common method to measure pain in mice, the tools described here, with the addition of a high-speed camera, are fully compatible with the setups that most labs currently use. As such, we do not foresee major hurdles in wide adoption of this methodology. To increase our fundamental understanding of the neurobiology of pain, and to translate our basic science findings in pain research to improved patient outcomes, the pain measurement tools in rodents must be robust. The platform described here should aid in that pursuit.

## METHODS

### Mouse strains

Mice for behavior testing were maintained in a barrier animal facility in the Carolyn Lynch or Translational Research Laboratory (TRL) buildings at the University of Pennsylvania. The Lynch and TRL vivariums are temperature controlled and maintained under a 12 hour light/dark cycle (7am/7pm) at 70 degrees Fahrenheit with ad lib access to food (Purina LabDiet 5001) and tap water. All procedures were conducted according to animal protocols approved by the university Institutional Animal Care and Use Committee (IACUC) and in accordance with National Institutes of Health (NIH) guidelines. C57BL6/J, A/J, 129S1/SvlmJ, Balb/cJ, DBA/1J, AKR/J, CBA/J, SJL/J, and TRAP2 mice (Fos-FOS-2A-iCre-ERT2) mice were all purchased from Jackson Laboratories.

### High speed imaging

Mouse behaviors were recorded at 2000 frames per second (fps) with a high-speed camera (Photron FastCAM Mini AX 50 170K-M-32GB - Monochrome 170K with 32GB memory) and attached lens (Zeiss 2/100M ZF.2-mount). Mice performed behavior in rectangular plexiglass chambers on an elevated mesh platform. The camera was placed at a ∼45° angle at ∼1-2 feet away from the Plexiglas holding chambers on a tripod with geared head for Photron AX 50. CMVision IP65 infrared lights that mice cannot detect were used to adequately illuminate the paw for subsequent tracking in ProAnalyst. All data were collected on a Dell laptop computer with Photron FastCAM Analysis software.

### Somatosensory behavior assays

Mice were habituated for a minimum of 5 days, for one hour each day, in the Plexiglas holding chambers before testing commenced. During baseline, mice were tested in groups of five and chambers were placed in a row with barriers preventing mice from seeing each other. On testing day, mice were habituated for an additional ∼10 minutes before stimulation and tested one at a time. Stimuli were applied through the mesh to the hind paw proximal to the camera. Testing only occurred when the camera’s view of the paw was unobstructed. Mice only received one stimulus on a given testing day (cs, db, lp, or hp) and were given at least 24 hours between each stimulus session. Stimuli were tested from least painful to most: cotton swab, dynamic brush, light pinprick, heavy pinprick. Cotton swab tests consisted of contact between the cotton Q-tip and the hind paw until paw withdrawal. Dynamic brush tests were performed by wiping a concealer makeup brush (e.l.f.TM, purchased at the CVS) across the hind paw from back to front. Light pin prick tests were performed by touching a pin (Austerlitz Insect Pins®) to the hind paw of the mouse. The pin was withdrawn as soon as contact was observed. Heavy pinprick tests were performed by sharply pressing this pin into the paw so that it was pushed upwards, without the breaking the skin barrier. The pin was withdrawn as soon as approximately 1/3 of the pin’s length had passed through the mesh.

### Automated paw tracking

We used ProAnalyst software to automatically track hind paw movements following stimulus application. This software allowed us to integrate automated and manually scored data, possible through the ‘interpolation’ feature within ProAnalyst. We were able to define specific regions of interest (paw), track, and generate data containing ‘x’ and ‘y’ coordinates of the paw through time, as well as velocity, speed, and acceleration information. In a subset of videos, additional manual annotation was performed for increased accuracy. For deep learning-based paw tracking with the Social LEAP algorithm, we pseudo-randomly chose a small set of training frames from each video and hand-labelled the paw and toe. We trained Social Leap to predict toe and paw positions in unlabeled video frames (>∼90% of total video frames). To generate trajectories, we overlaid the inferred x, y positions of the toe and paw in each video frame on a single still image corresponding to the apex of the mouse’s paw during the assay.

### Development of PAWS software to quantify pain behaviors

Behavioral features were extracted from raw paw position time series in an automated and standardized procedure. First, the start and end of paw movement (paw at rest on the ground) were identified, and analysis was restricted to this time window. Peaks in paw height were then determined based on Savitsky-Golay smoothed estimates of paw velocity, and the first peak identified. The time of the first peak (designated t*) was used to separate pre-peak behavioral feature calculations from post-peak calculations. To differentiate shaking from guarding in the post-peak period, we constructed a moving reference frame based on the principal axis of paw displacement across a sliding window (0.04 seconds in duration) for each time point, and identified periods of consecutive displacements above a specified threshold (35% of maximum paw height) as periods of shaking. Note that in the construction of the moving reference frame the principal axes of variation were recovered via PCA, which is not invariant to the sign of the recovered axes. Since displacement is measured over time it is sensitive to reversals in sign along the axis we measure it. We therefore ensured consistency by using the axis direction minimizing the angular deviation from the axis recovered at the previous time step.

### Automated behavioral feature selection method using SVD

As a point of comparison for the behavioral measures quantified as pre- and post-peak paw features using common measures from the literature, we employed a simple, unsupervised machine learning technique to generate candidate behavioral features based directly on variation in the paw trajectory timeseries themselves. Time series were aligned by the time of the first peak in paw height, t*, resulting in time series of 4000 discrete measurements per paw dimension (sampled at 2000fps). This representation was then compressed by a factor of 87.5% by retaining only the first 500 coefficients per spatial dimension of its discrete cosine transform (DCT-II), which nonetheless maintained paw position accuracy to within 1.5 and 0.11cm RMSE in x (anterior/posterior) and y (height), respectively. A singular value decomposition of this compressed description (both x and y dimensions combined per time series) was used to load the correlated variation within time series onto a small number of dimensions ordered by decreasing variation capture. The number of SVD dimensions ultimately retained, 23, was determined by choosing the dimensionality that maximized the LDA probability correct under strain leave-one-out cross-validation.

### Drugs

4-hydroxytamoxifen (Hello Bio, #HB2508) prepared in Kolliphor EL (Sigma, #27963), Clozapine-N-oxide (Hello Bio, #HB6149), and 0.9% sodium chloride (Sigma, #S3014).

### Viral Reagents

For chemogenetic manipulation of BLA pain-active neurons, we intracranially injected 200 nL of AAV5-*hSyn*-*DIO*-*h*M3D(G_q_)- *mCherry* (Addgene, titer: 7 × 10^12^) into both the left and right BLA at coordinates AP: -1.4 mm, ML: ±3.1 mm, DV: -4.2.

### Stereotactic injections and surgical procedures

We conducted surgeries under aspetic conditions using a small stereotaxic instrument (World Precision Instruments). We anesthetized mice with isofluorane (5% induction, 1-2% maintenance) during the entire surgery and maintained body temperature using a heating block. We injected mice with a beveled 33G needle attached to a 10 µL syringe (Nanofil, WPI) for delivery of 200 nL of viral reagent at a rate of 40 nL/min. After viral injection, the needle remained at the injection depth for 10 minutes before slow withdrawal over 2 minutes. After surgery, we maintained the animal’s body temperature using a heating pad.

### Targeted recombination in active populations (TRAP) of BLA pain ensemble

Transgenic female TRAP2 mice were injected with an AAV at P46-73. Two weeks after injection, mice were stimulated with a noxious pin prick on the left hind paw every 30 seconds for 10 minutes. One hour later, mice were injected with 4-hydroxytamoxifen (4-OHT) to induce genetic recombination. Eight weeks following 4-OHT administration, behavior of mice was examined. Mice were sacrificed via transcardial perfusion 4 months after viral injections. Brains were collected and sectioned at 50 µm on a cryostat. Tissue was mounted and imaged on a fluorescent Keyence microscope.

